# Patch characteristics and domestic dogs differentially affect carnivore space use in fragmented landscapes in Southern Chile

**DOI:** 10.1101/2020.12.20.423635

**Authors:** Rumaan Malhotra, Jaime E. Jiménez, Nyeema C. Harris

**Affiliations:** Applied Wildlife Ecology Lab, Ecology and Evolutionary Biology, University of Michigan, 1101 N. University Ave, Ann Arbor, Michigan 48106; Advanced Environmental Research Institute, Department of Biological Sciences, University of North Texas 1155 Union Cir, Denton, Texas 76203

**Keywords:** camera survey, *Canis lupus familiaris*, disturbance, fragmentation, invasive species, landscape, Lycalopex, occupancy

## Abstract

In an increasingly anthropogenic world, native species face multiple interacting threats. Habitat fragmentation and domestic dogs are two such perturbations threatening terrestrial mammals globally. Here, we implemented a camera trap survey in the fragmented central valley/Andean foothills transition of the Los Lagos Region in Southern Chile to evaluate space use of native carnivores in a landscape comprised of patches of native forest amidst a matrix of pastureland. Using an occupancy modeling framework to account for imperfect detection, we examined the impacts of dogs and landscape metrics of fragmentation on three mesocarnivores – the foxes culpeo (*Lycalopex culpaeus*) and chilla (*Lycalopex griseus*) and the wild cat güiña (*Leopardus guigna*). Factors driving occupancy differed for each of the native species, while detection rates for both canid species increased with dog occupancy. We found that a small (12%) simulated increase in dog occupancy negatively impacted the spatial use of the culpeo. Habitat loss and fragmentation were positive drivers for the chilla and the dog, and indirectly impacted the culpeo through the domestic dog. The güiña did not respond to fragmentation and other habitat covariates or dog occupancy. Instead, all native carnivore species temporally partitioned diel activity with dogs. We highlight that the effects of dogs or fragmentation are not ubiquitous across the carnivore guild with varied tolerance. However, future conditions of increased fragmentation and habitat loss will likely increase the potential contact between domestic dogs and native carnivores.

## 1. Introduction

Fragmentation and habitat loss remains a global threat to biodiversity, increasing isolation between suitable habitat with emergent consequences from edge effects (Haddad et al. 2015; Pfeifer et al. 2017). These physical changes to landscapes also impact abiotic factors, altering radiation fluxes, wind patterns, and the hydrological cycle to increase heterogeneity within and across habitats (Saunders et al. 1991; Schmidt et al. 2017). Globally, 70% of forests are within 1 kilometer of an edge and are becoming increasingly fragmented resulting in abundances for over 85% of vertebrates being impacted by edge effects (Haddad et al. 2015; Pfeifer et al. 2017; Montibeller et al. 2020). The negative effects of fragmentation remain highly debated given idiosyncratic impacts across species and ecological interactions (Fahrig 2013; Rielly-Carroll & Freestone 2017; Fletcher et al. 2018; Fahrig et al. 2019; Harrison & Banks□Leite 2020). While species may be able to inhabit edge habitats, they may be excluded via biotic factors such as competition or predation (Michel et al. 2016). Additionally, fragmentation may interact with other factors such as habitat loss, fire prevalence, and hunting, exacerbating impacts or making it difficult to ascertain the individual drivers that alter species or interactions (Cochrane 2001; Peres 2001; Bartlett et al. 2016).

Domestic dogs (*Canis lupus familiaris*) represent another global threat to biodiversity as the most abundant carnivore worldwide with a global population estimated at 700 million (Hughes & Macdonald 2013). Therefore, dogs are a widespread invasive species that can commonly exploit fragmented landscapes as they more easily permeate from areas of human residence (Oehler & Litvaitis 1996; Broadbent et al. 2008; Paschoal et al. 2018). Dogs commonly harass and kill native carnivores, compete for prey species, and transmit pathogens to wild populations, threatening native carnivores (Laurenson et al. 1998; Vanak & Gompper 2009; Doherty et al. 2017). These disturbances can alter activity patterns and reduce relative abundance of native carnivores. For example, native carnivores in Madagascar exhibited spatial avoidance when domestic dogs were present, and were more likely to be replaced by dogs in degraded forests near human settlement (Farris et al. 2016, 2017). Similarly, chilla fox (*Lycalopex griseus*) visits to scent stations in Southern Chile were negatively correlated with dog presence, and telemetry data showed that foxes rested in a habitat type that was not preferred by dogs (Silva-Rodríguez et al. 2010a). In general, how dog-wildlife interactions are facilitated by habitat destruction is largely unstudied, and it is unknown whether habitat destruction and dogs have similar or opposing impacts on native carnivores, or work in concert. Given the pervasiveness of both dogs and fragmentation as major disturbances to native species, it is surprising that few studies measure and compare the synergistic effects of both.

The susceptibility to dog harassment/killing is largely size biased and thus, intensified for mesocarnivores. However, the impacts of fragmentation on carnivores is harder to predict because many aspects of their ecology such as prey availability, habitat quality that are also impacted. For example, a disturbance from fragmentations shuffles species distributions and facilitates the invasion of nonnative competitors or other domestic species (Crooks 2002; Echeverría et al. 2007; Jessen et al. 2018). Mammals vary in their sensitivity to fragmentation and in their adaptive responses from fragmentation (Crooks 2002; Janecka et al. 2016; Smith et al. 2019; Palmeirim et al. 2020). Large-bodied mammalian carnivores are particularly susceptible to fragmentation and edge effects due to their relatively small population sizes, slow growth rates and extended habitat requirements and corresponding home ranges (Schipper et al. 2008). The coupling of natural history characteristics, landscape structure, and anthropogenic pressures from human persecution result in carnivores, of all sizes, being among the most threatened groups of mammals worldwide (Karanth & Chellam 2009; Estes et al. 2011; Ripple et al. 2014). However, the impacts of fragmentation are less clear for mesocarnivores, many of which are generalists and have smaller home ranges than their larger counterparts, and thus may be more resistant to or even benefit from fragmentation (but see Crooks et al. 2017; Rocha et al. 2020). For example, Massara et al. (2016) found that the occupancy of generalist mesocarnivores was negatively tied to reserve size throughout the remnant patches of the Atlantic Forest in Brazil,.

Land owned privately by smallholders has been largely omitted from studies on native carnivores and interactions between the carnivore guild and human pressures. These understudied areas represent a later stage of the fragmentation process; rather than a contiguous protected area with an edge riddled with encroaching pastures and other human use Given the increased anthropogenic impacts in these areas, it is likely there is increased dog presence as well (Paschoal et al. 2018). These ‘working’ lands have traditionally been discounted in their value for conservation, being considered largely as the hostile matrix that native species must navigate between protected areas. However, recent findings show that patches within these agriculture-dominated lands can hold considerable biodiversity and can have high conservation value (Kremen & Merenlender 2018; Lindenmayer 2019; Wintle et al. 2019). Given the huge potential of agricultural lands for conservation, comprising a third of the ice-free land on the planet (Ramankutty et al. 2018), there is a increasing recognition that co-production of science with private landowners on working lands is necessary and perhaps unavoidable in efforts to maintain biodiversity (Naugle et al. 2020).

In the Valdivian temperate forests biodiversity hotspot of Chile, both fragmentation and the presence of domestic dogs are widespread and potentially devastating endemic species (Myers et al. 2000). These forests are being rapidly lost and converted to exotic plantations and pasturelands (Echeverría et al. 2008; Echeverría et al. 2012). The current conservation estate is insufficient in meeting goals to maintain the biodiversity value of these forests because protected areas are restricted to the inland Andes rather than the endemic-rich coastal areas (Smith-Ramírez 2004). The central valley, which formerly connected the coastal and montane sections as contiguous forest, has been heavily deforested and now dominated by cow pastures and exotic plantations. Today, only small patches of native forests remain as available wildlife habitat that are interspersed throughout this landscape that are privately-owned and managed (Figure 1). Free-ranging domestic dogs pose a major threat to the persistence of at least two mammal species of conservation concern, pudu (*Pudu puda*, IUCN status of Vulnerable) and Darwin’s fox (*Lycalopex fulvipes*, IUCN status of Endangered) (Silva-Rodríguez et al. 2010b, 2016).

**Figure 1.**
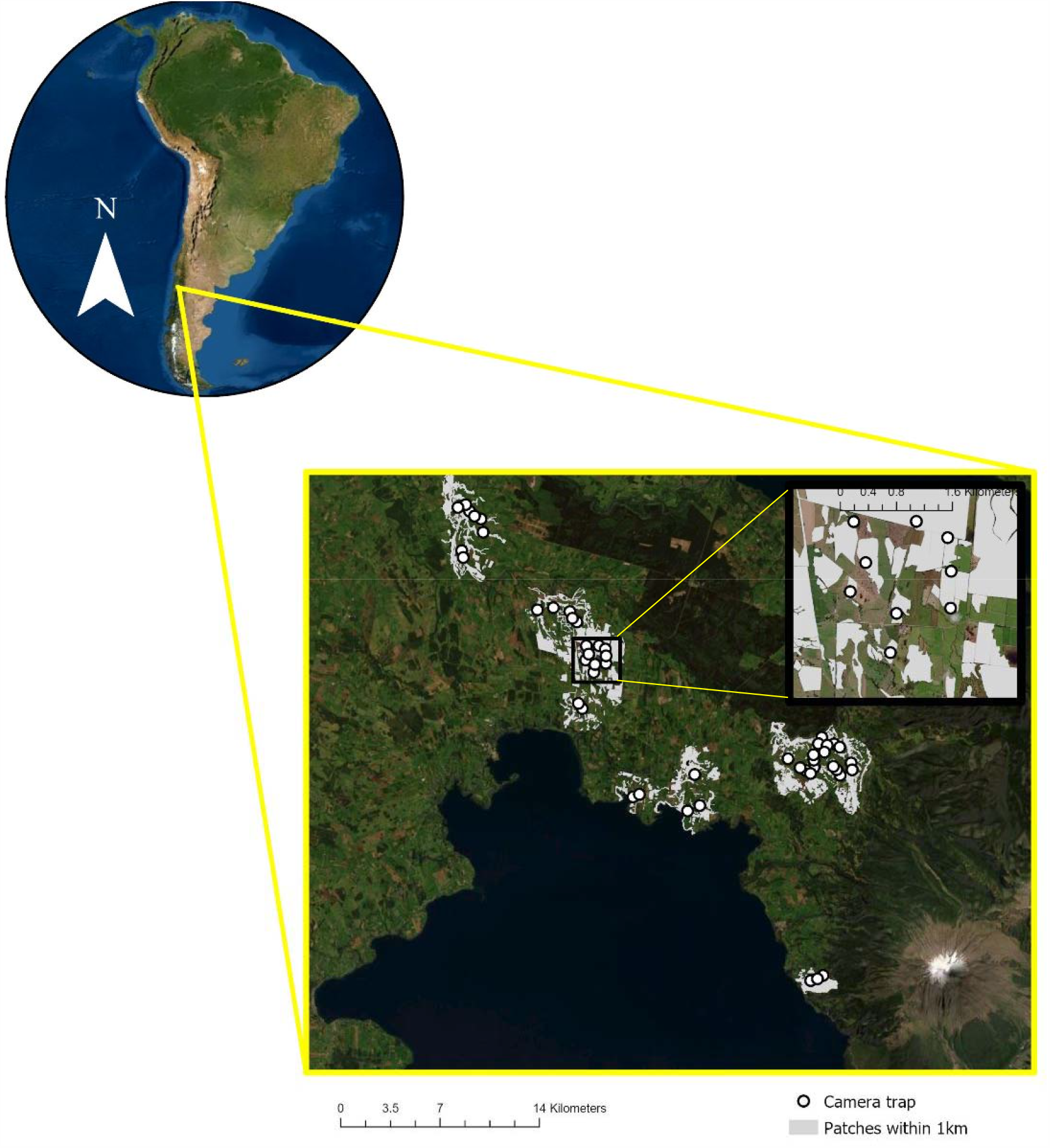
a) Study area located in the Los Lagos Region of south-eastern Chile. b) Landscape level distribution of camera deployment throughout patches of native forest straddling the Volcano Osorno.

Here, we determine the relative consequence of fragmentation, the presence of dogs, and the interaction between the two on the spatial use of native carnivores. Specifically, we surveyed privately-owned forest patches that were outside of protected areas or forestry company ownership using remotely-triggered cameras. We expected fragmentation metrics to be more important than dog space use in explaining the occupancy of forest specialists (e.g., güiña, *Leopardus guigna*). In contrast, we also expected that in these largely altered landscapes dog occupancy would be the major driver of native canid spatial use, due to the immediate threat they present, and induced ‘fear effects’ (Palomares & Caro 1999; Vanak et al. 2009; Vanak & Gompper 2010). We hypothesized that increasing patch isolation and reducing the proportion of forest would be important drivers of dog occupancy, providing evidence that their presence is facilitated by fragmentation (Figure 2). Our work will enhance our understanding of native carnivore occurrence in the later stages of human-altered landscapes and reconcile the relative contributions of interacting threats from fragmentation and domestic dog presence.

**Figure 2.**
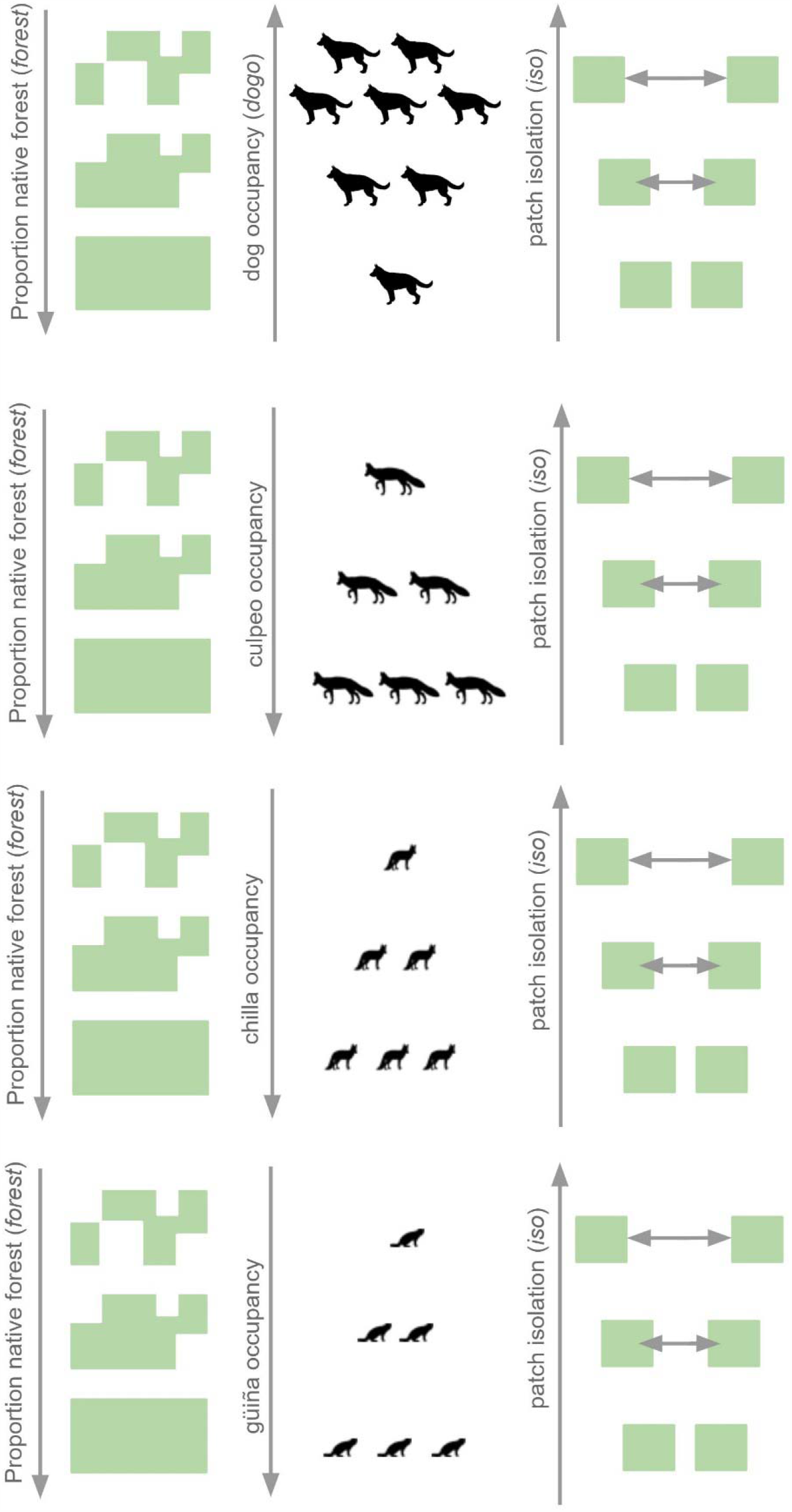
Hypothesized facilitation of dog occupancy by habitat loss and fragmentation with expectation that decreasing proportion of native forest and increasing patch isolation would promote higher dog occupancy. Expectations for native carnivore response to fragmentation were opposite those of domestic dogs, with native carnivore occupancy expected to decrease with decreasing forest and increasing patch isolation.

## 2. Methods

### 2.1 Study area

We surveyed the carnivore community in the Los Lagos region of Chile, near the city of Osorno, between Lago Rupanco and Lago Llanquihue (40° 76′ to 41° 21′ S, 72° 54′ to 72° 97′ W, Figure 1). This area is characterized by Valdivian temperate rain forest (mean temperature 3-23°C) with a cold, rainy winter season between May and September (1346 mm annual rainfall) and mild summers (en.climate-data.org). The landscape, formerly dominated by native forest, is currently dominated by pastures that are used primarily for cattle as well as plantations of pine (*Pinus radiata*) and eucalyptus (*Eucalyptus* sp.) with small stands of native forest. The study area is relatively flat and sandwiched between a large protected area (Parque Nacional Vicente Rosales) on the eastern edge and the Osorno metropolitan area on the western edge. Native forest patches were mostly made up of degraded strips along the edges of pastures comprised of a mix of *Lophozonia obliqua, Nothofagus dombeyi*., *Persea lingue*, and *Laurelia sempervirens* with a bamboo understory (*Chusquea quila*).

### 2.2 Camera trap survey

We deployed 50 remotely-triggered cameras (Reconyx© PC 850, 850C, 900, 900C) in native forest patches throughout the study area from June to August 2019, during the austral winter. We affixed cameras to trees (minimum diameter 0.25 m) with cable locks and placed 0.5 m off the ground. We used signs of animal activity such as game trails and scat to determine the specific micro-site location of camera placement, to maximize detections. Cameras were placed at least 0.5 km apart from each other, and efforts were made to place within the core of each patch if minimum spacing allowed. Each camera was baited with canned mackerel placed inside a perforated bottle, wired down to keep animals from accessing or removing the bait. Camera settings included: high sensitivity, one-second lapse between three pictures in a trigger, rapidfire (no quiet period between triggers).

At the end of the survey period, images were retrieved from the cameras and identified by a single observer (R. Malhotra) to the species level. After image identification, we applied a 30-minute quiet period to ensure independence of species detections (Wang et al. 2015; Suraci et al. 2016). These images and the associated site-level environmental variables (explained below) were used to estimate individual species occupancy. We used Moran’s I in ArcPro (vers. 2.3.1) and did not find evidence of spatial autocorrelation. The ‘camtrapR’ package in Program R was used to organize camera trap images and extract data for modeling.

### 2.3 Occupancy modeling

Using single-species single-season occupancy models (MacKenzie et al. 2003), we evaluated the impacts of habitat degradation on the occupancy (Ψ) and detectability (*p*) of dogs, and evaluated the impacts of habitat degradation and dog occupancy on the occupancy and detectability of three focal native species: the chilla (*Lycalopex griseus)*, culpeo (*Lycalopex fulvipes*), and güiña (*Leopardus guigna*) (Figure 3). We expected that increasing habitat loss (*forest*) and patch isolation (*iso*) would reduce native carnivore occupancy, and that native species occupancy would be inversely related to dog occupancy (*dogo*). We first separated species detections into 7-day observation periods. We then modeled detection probabilities for each species holding occupancy constant, and then used the best detection models to model the occupancy for each species.

**Figure 3.**
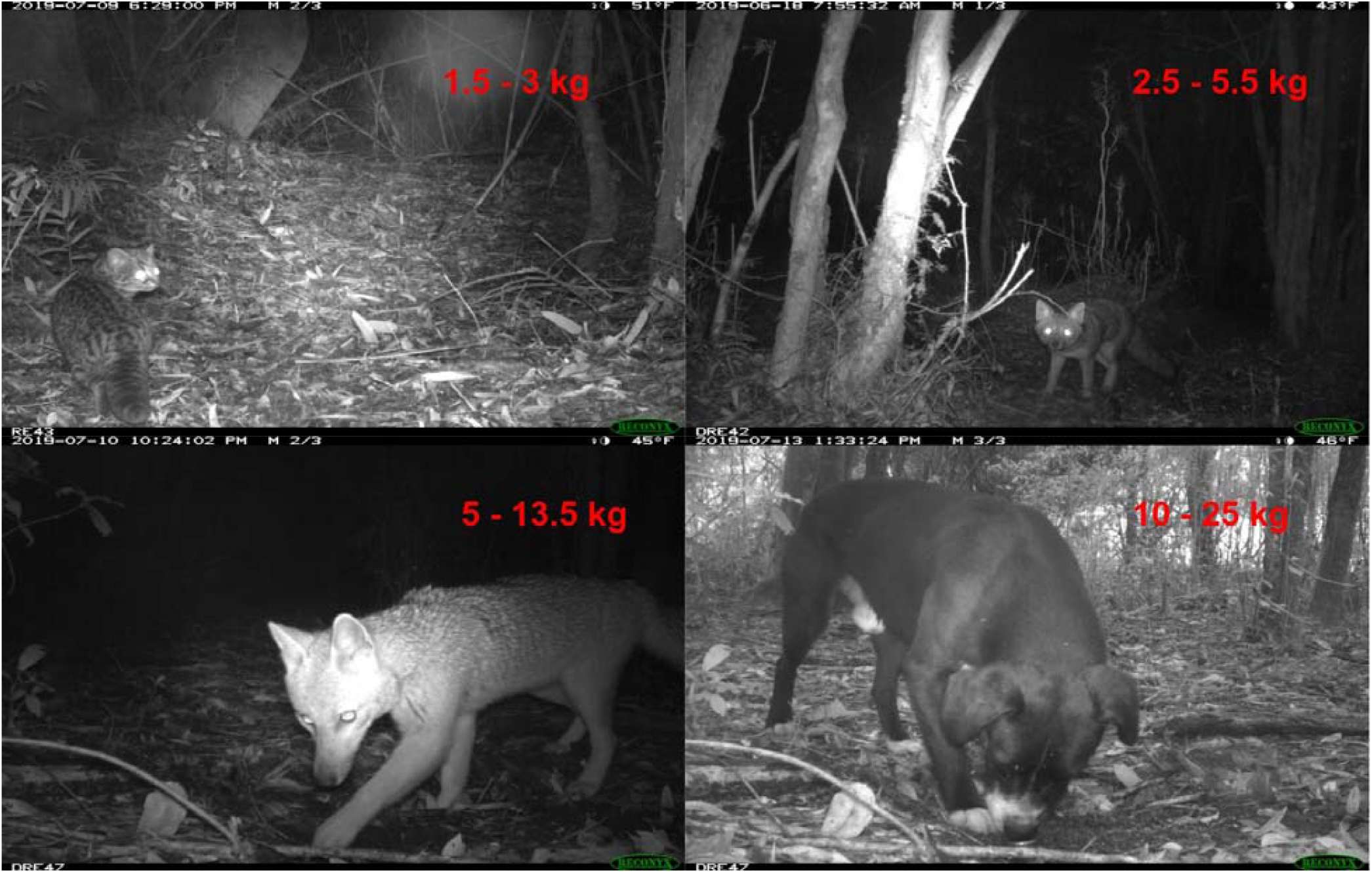
Focal carnivores in study for size comparison of the three native species relative to and domestic dogs: the güiña, chilla, domestic dog, and culpeo from top left clockwise. Note that the upper weight range of the culpeo likely represents more southern parts of the range than the study area; they are relatively bigger than chillas, and smaller than dogs. Photo credit: Applied Wildlife Ecology Lab.

#### 2.3.1 Detection covariates

We modeled detection probabilities with covariates hypothesized to influence visibility of species on camera images. We measured understory cover using a point-intercept method, with the height measured every meter for 10m in the four cardinal directions surrounding each camera (Karl et al. 2017). We then aggregated values for the understory cover into three categories: 0m (no understory), 0.01-0.25m, and 0.26-0.5m. Understory at 10m (*10uds*) is an average of all understory measurements taken every meter within a 10-meter radius of the camera tree (40 measurements per camera site). We expected the understory covariate to have a negative relationship with detectability of all species, as animals passing by would be obscured by increased understory towards the height of camera placement. Understory was not included in the detection model for the culpeo, as inclusion prevented convergence of the global model. We first modeled occupancy of dogs using habitat covariates (*10uds, forest, iso, sm*), and then included the resulting site level estimates as the *dogo* covariate for native species models (Figure 3). Patches were digitized in ArcPro (vers. 2.3.1) using high resolution satellite imagery from 2018 (Maxar Vivid) to obtain proportion native forest cover (*forest*), patch isolation (*iso*) and a metric of edge effects (*edge*) estimates. Patch isolation (*iso*) was measured as the mean border-to-border distance to the nearest patch within a 1-km radius of each camera. The *edge* covariate was measured as the mean ratio of patch perimeter size to patch area for all patches within a 1-km radius of each camera. However, the edge covariate was excluded from the final global model, as it was highly correlated with both *forest* and *iso* covariates (*p* < 0.01). Both *iso* and f*orest* covariates were expected to have a positive relationship with native species occupancy.

We estimated relative abundance of small mammals (*sm*), as a metric of prey availability, using the total number of all independent lagomorph, rodent, and shrew opossum triggers per camera standardized by the number of trap nights. Camera type (*cam*) was included to distinguish between white-flash cameras and infrared cameras with the expectation that white-flash cameras would have lower species detection despite better nighttime image quality, due to possibly startling species. Lastly, trap nights (*trap*), the number of nights an individual camera was operational to collect species detections, were included to determine if sampling effort affected detection rates. However, we expected no effect of trap nights, given similar sampling periods for all the cameras.

#### 2.3.2 Occupancy covariates

Occupancy for each species was modeled with dog occupancy (*dogo*) and habitat covariates (*10uds, forest, iso, sm);* edge was highly correlated with *forest* and omitted from the model. To test whether the impact of dogs on native carnivores was facilitated by lack of understory, we included an interaction term between dog occupancy and average patch understory height within 10m (*dogo*\**10uds*). Individually, we expected dog occupancy to reduce occupancy of all native species, and for understory height to increase the occupancy of the smaller native carnivores (güiña and chilla). As an interaction term, we expected increasing understory to decrease the effect of dog occupancy on native species occupancy. We expected higher prey availability to be a positive driver of native species occupancy, but unimportant for the domestic dog, which in this area would be classified as ‘free-ranging’ by Vanak & Gompper (2010) and thus would rely on human food subsidies.

#### 2.3.3 Model evaluation

The dog global model included mean understory height within 10m (*10uds*) and camera type and trap night (*cam, trap*) covariates for detection, and understory (*10uds*), prey (*sm*), and fragmentation (*forest, iso*) for occupancy. Native species global models used the same covariates as the dog model, with the addition of dog occupancy (*dogo*) for detection, and an interaction term for dog occupancy and understory (*dogo*\**10uds*) for occupancy. All detection and occupancy covariates were tested for correlation by site using Pearson’s R. Model ranking was carried out using Akaike Information Criterion, corrected for small sample sizes (AICc), or quasi-AICc (QAICc) if the global model was overdispersed (c-hat > 1.2), with the top model being defined as the one with the lowest AICc or QAICc score. Goodness of fit was tested for all top models (<2 Δ AICcs or QAICcs of the highest rank model) using a Chi-square statistic. All occupancy modeling was completed in the ‘unmarked*’* package in Program R vers. 3.6.2.

#### 2.3.4. Threshold response to dogs

We interpreted the β coefficient of *dogo* and confidence intervals overlapping zero when occurring in top models to conclude significant effects of dogs on native species occupancy. When the top models included *dogo* as a covariate with a non-significant negative coefficient, we determined the threshold level of dog occupancy required for *dogo* to become a significant negative driver on native species occupancy. We incrementally increased the value of the *dogo* to the maximum occupancy value (1), a single camera at a time. The order was determined by ranking cameras from highest to lowest *dogo* value.

### 2.4 Temporal use

As sympatric carnivores may be more likely exhibit temporal instead of spatial partitioning to promote coexistence, we estimated pairwise temporal overlaps for all species, and compared the overlap of native species pairs with the overlap of native species-dog pairs. We expected native species to have nocturnal activity patterns, dogs to have a diurnal activity pattern, and for native species to overlap more with the other native species than with domestic dogs. We plotted the temporal activity distributions of each species and determined the degree of overlap between pairs (Δ) with 95% confidence intervals generated by 10,000 parametric bootstrap iterations. Δ values range from 0 indicating completely distinct and non-overlapping temporal activity to 1 indicating complete overlap between the comparison groups. Δ_1_ was used for comparisons when one of the sample groups had less than 50 triggers; otherwise Δ_4_ was used to estimate temporal overlap between species pairs (Ridout & Linkie 2009). We then used the Mardia-Watson-Wheeler test to determine if the temporal patterns varied significantly between individual species, which compares two sets of circular data and determines if there is homogeneity in the means or variances. We implemented the temporal analyses using the ‘overlap*’* and ‘circular*’* packages in Program R.

## 3. Results

We detected all three native carnivore species over a total effort of 3500 trap-nights. Naïve occupancy estimates for the güiña (n=56 independent triggers), chilla (n = 225), and culpeo (n=39) were 0.51, 0.59, and 0.16 respectively. Domestic dogs were fairly common (n=64) found at 20/49 camera sites (naïve Ψ = 0.41). Additional native carnivores that were detected, although rare, included the chingue (*Conepatus chinga*, n=13) and the puma (*Puma concolor*, n=4). We also detected two additional non-native species: the mink (*Neovison vison*, n=20) and domestic cat (*Felis catus*, n=21). Darwin’s fox (*Lycalopex fulvipes*) was not detected during our camera survey in the area.

### 3.1 Detection of carnivores

Our study area was comprised of an understory that ranged from completely open to thickets of dense vegetation with specific camera sites comprising no understory to over two meters in height. As such, we expected detection to vary by understory, depending on species preference on microsite selection for dense vegetation, and the ability of understory to reduce the visibility range for a camera trap. For the chilla (β = −6.16, SE = 1.38) and dog (β = −7.44, SE = 1.84), *10uds* was a strong driver of detection probability, decreasing the detectability for both species (Table S1). For both chilla (β = 1.77, SE = 0.37) and culpeo (β = 3.23, SE = 0.834), *dogo* increased detectability. The null model best described güiña detection; that is, no effect of covariates improved model fit.

### 3.2 Occupancy of carnivores

Overall, modeling occupancy with covariates and accounting for imperfect detection improved our understanding of carnivore space use. Chilla foxes had the highest overall occupancy (Ψ = 0.67), while culpeos had the overall lowest occupancy, but nearly doubled from the naïve estimate (Ψ = 0.36). Güiña was the only species for which the null model was the best model, and the occupancy estimate was thus the same as the naïve estimate (Ψ = 0.51). In comparison to the native species, dog occupancy was higher than the culpeo and güiña, but lower than that of the chilla (Ψ = 0.58).

Factors driving occupancy of native carnivores varied by species (Figure 4, Table 2). Despite the importance of *10uds* for species detection, it did not appear in the best model for any species. It was however a negative driver of chilla occupancy in 4/10 top models which had comparable weight to the best model (Table 2). Given the reliance of mammalian carnivores on prey, unexpectedly, *sm* was important only for the occupancy of the culpeo (β = 1.05, SE = 0.53).

**Figure 4.**
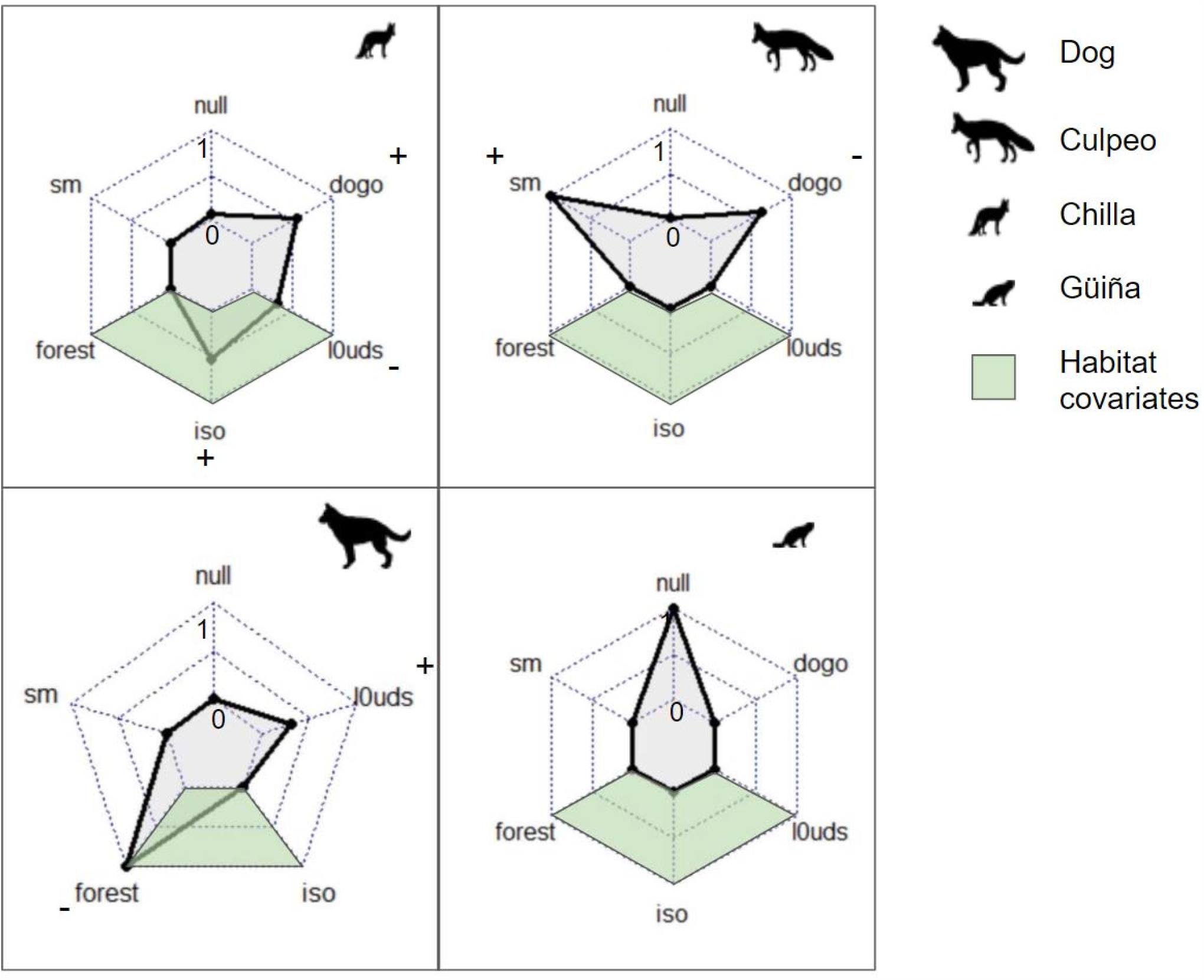
Relative importance of each covariate on species occupancy based on summed model weights for top model sets (< 2 ΔAIC/QAIC). Positive (+) and negative (–) signs correspond to the direction of beta coefficients from each model set, and were consistent within top model sets.

We aimed to contrast the ecological consequences of habitat destruction (loss and fragmentation), and dog occurrence on the space use of native carnivores (Table 2). Habitat metrics were important drivers of chilla and dog occupancy but did not appear in the model sets for culpeo or güiña. For example, *iso* was in the top two models for the chilla (β = 0.09, SE = 0.04), and was positively correlated with occupancy. *Forest* did not appear to be important for occupancy of any native species. We found that *forest* was however important for dogs (β = −26.06, SE = 12.1), with increasing proportion of native forest decreasing dog occupancy. *Dogo* appeared in 4/5 top models for culpeo (including the best model) and was important for model fit for the culpeo, but was not a significant driver of culpeo occupancy (β = −4.19, SE = 2.74).

Similarly, results varied in quantifying responses of native carnivore occupancy to domestic dog presence. For chilla, *dogo* was not in the best chilla model but appeared as a positive driver in 6/10 top models, which had comparable weight to the best model (Table S1). The *dogo* covariate was not influential, positive or negative, on occupancy for güiña. Culpeo was the only species with *dogo* in the top model with a negative (non-significant) βcoefficient. The dog landscape level occupancy from the top model was 0.58. Increasing dog occupancy to 0.65 (an increase of 12.1%) resulted in *dogo* becoming a significant negative driver of culpeo occupancy (Figure S2).

Ultimately, we rejected our hypothesis of dog effects on native carnivores being facilitated by lack of understory. We found no evidence for the interaction of *dogo* and *10uds* affecting occupancy for native mesocarnivore species occupancy. Overall, we conclude that landscape characteristics via metrics of increasing fragmentation have similar positive effects for both generalist native carnivores and for domestic dogs (Figure 4).

### 3.4 Temporal activity

We evaluated temporal activity patterns of all of our study species to determine if there was evidence for temporal avoidance with domestic dogs. Activity patterns for the three native carnivores was largely restricted to the nocturnal and crepuscular hours. Overlap among the native carnivores was high (Δ ranging 0.78 – 0.89) and did not vary significantly among pairs (Table S1: *p values*: 0.08-0.79). In contrast, domestic dog activity was almost entirely diurnal, resulting in significantly different activity patterns from native species (Δ ranging 0.35-0.43; *p* < 0.001). Furthermore, 95% confidence intervals for Δ dog-native species pairs and for Δ native-native species pairs did not overlap in a single case, indicating that native species overlapped significantly more with other native species than they did with dogs.

## 4. Discussion

The threats that mammals face from habitat loss and fragmentation are especially relevant in the context of the temperate rainforests of central Chile, which have included rapid deforestation and fragmentation in the past 50 years (Echeverría et al. 2006; Nahuelhual et al. 2012; Uribe et al. 2020). An additional human-related threat is the presence of domestic dogs, which antagonize native species and preferentially use the pasturelands that separate the remaining native forest patches (Silva-Rodríguez et al. 2010a). Our results from the remnant forest patches within an agricultural matrix in southern/central Chile indicate that both fragmentation and domestic dogs have differing effects on native carnivore occupancy.

The effects of habitat loss and fragmentation for native carnivores are important to explore given future trends in the deforestation of the region. Our study site represents a landscape that has already undergone extensive habitat loss and fragmentation; primarily agricultural land with only remnant patches of native forest. However, even in already heavily degraded regions such as our study site, trends indicate that native forest throughout Chile continues to decline, with available habitat patches decreasing in size (Echeverría et al. 2008; Echeverría et al. 2012; Miranda et al. 2015). For the species included in this study, at first glance our results suggest that this landscape degradation does not pose an immediate concern. For the chilla, the positive correlation between patch isolation and occupancy is likely a reflection of the ecology of the fox, which primarily forages in the open fields that comprise the matrix between forest patches (Silva-Rodríguez et al. 2010a). However, this species also utilizes interior habitat of these native forest patches as a refuge, and thus, would likely have negative consequences if forest patches fell below a threshold sizes (Silva-Rodríguez et al. 2010a). Our results for culpeos and güiñas, which did not show any response to either habitat loss or patch isolation could indicate that: a) these species are plastic in their habitat requirements; b) fragmentation and habitat loss have not reached a sufficient threshold to elicit a response; c) there is a time lagged ‘extinction debt’, or d) these species are tracking spatial patterns of prey, predator, or competitor species instead (Hanski & Ovaskainen 2002; Ryall & Fahrig 2006; Swift & Hannon 2010; Halley et al. 2016). The model results for the culpeo seemed to suggest this latter mechanism, as they were positively driven by prey availability and dog occupancy was consistent in the top models having a negative coefficient (though note that it was not significant). While landscape characteristics did not appear as a negative driver in any native species models, the inverse relationship between dog occupancy and proportion of native forest means that as habitat loss increases in this region, it will likely mean less refuge habitat for native species, and higher exposure to domestic dogs (Torres & Prado 2010; Paschoal et al. 2018).

We expected dogs to influence native carnivore occupancy because of their documented impact on small carnivores through interference and exploitation competition, and the increased mortality risk they pose as disease reservoirs and vectors (Laurenson et al. 1998; Rhodes et al. 1998; Sillero-Zubiri et al. 2004; Vanak & Gompper 2009, 2010). Dogs are potential competitors with native carnivores and have been linked to the decline of the native pudu, a potential prey item for the two fox species in this study (Silva-Rodríguez & Sieving 2012). Despite the threat that a dog encounter presents, dog occupancy did not clearly present a negative driver of native species occupancy, and only featured as a non-significant negative covariate for culpeo top models. While this partially fit our expectation that native canids would more likely have antagonistic interactions with dogs and exhibit avoidance, we expected the smaller chilla fox to be more susceptible and affected (Donadio & Buskirk 2006; Vanak & Gompper 2009). Previous studies corroborate this expectation as dogs enforce interference competition to alter space use and have been observed harassing and killing chilla (Silva-Rodríguez et al. 2010a). A lack of a negative response from chillas to dogs using an occupancy framework could indicate that foxes were avoiding dogs at finer spatial or temporal scales, or that dog density was not sufficiently high to elicit a spatial avoidance (Zapata-Ríos & Branch 2018; Qi et al. 2020). Indeed our analysis of activity patterns suggests temporal partitioning as a mechanism for avoidance of dogs (Kronfeld-Schor & Dayan 2003; Schuette et al. 2013). In contrast to chillas, culpeos did indicate a potential response to dogs at a landscape level in congruence with recent work in the Andes (Zapata-Ríos & Branch 2018). Despite the differences in the landscape histories with our study conducted in a historically contiguous forest, while the Zapata-Ríos & Branch (2018) study occurred in the historically patchy Ecuadorian Andes, we both found that culpeos could respond to dogs rather than to habitat loss and fragmentation. Congruent with their dog occupancy estimates (Ψ-=0.66, range: 0.53 – 0.73), a projected 12% increase in dog occupancy in our study site for it to significantly reduce culpeo occupancy.

While dogs had opposite effects on the occupancies of the native fox species, they increased detection for both the culpeo and the chilla. Movement data for canids highlight quicker speeds through riskier areas, which would likely impact detection rates (Péron et al. 2017; Broadley et al. 2019). Thus, increased detection for the fox species may reflect a finer scale response to the risk posed by domestic dogs, rather than a broader change in spatial use (Broekhuis et al. 2013).

Fragmentation can facilitate the spread of invasive species through numerous pathways, such as roads increasing the occurrence of dogs (Loss et al. 2013; Moreira-Arce et al. 2015). Yet, few occupancy studies have looked at the impacts of both dogs and habitat loss and fragmentation on native carnivores. Our dog occupancy model revealed that dog occupancy decreased with proportion of native forest, providing evidence for the interaction between deforestation and dogs. Whether this interaction impacts native carnivores can be intuitively answered when we see that dog occupancy can be a negative driver of culpeo occupancy when surpassing a threshold. In ‘working’ landscapes this is particularly relevant as habitat loss and dog occupancy will likely continue to increase over time. Our occupancy results suggest that the spatial use of both native fox species (indirectly in the case of the culpeo, through dog occupancy) is tied to fragmentation and habitat loss. Furthermore, this change in the landscape increases the exposure of both native foxes to the threat of a dog encounter (Farris et al. 2017, 2020). In the currently remaining native forest stands that we surveyed, the largely nocturnal temporal use of native species provides a likely avoidance mechanism (Gerber et al. 2012; Shores et al. 2019). This temporal avoidance mechanism may be particularly important for generalist species such as the chilla, which our occupancy models show is similarly benefitted by habitat degradation as dogs. Future studies that investigate fragmentation on antagonistic interactions would prove beneficial to determining impacts on carnivore community structure (Magrach et al. 2014).

Our study gives us insight into the drivers of native carnivore space use in ‘working’ landscapes rather than the protected areas that represent ideal and untouched habitats. By situating our study on privately-owned smallholder lands, we also have the unique opportunity to inform the conservation of species in these increasingly anthropogenic landscapes. Many landowners do not have access to camera traps, and thus are unlikely to encounter elusive carnivores that are present even in small patches of native forest along the edges of their pasturelands. While voluntary strategies have greater social acceptance, they are not possible without landowners first having the knowledge of what species are on their land (Kamal et al. 2015). By partnering with landowners, scientists and managers can facilitate species conservation in these important landscapes which are not typically considered conservation targets (Naugle et al. 2020).

## Acknowledgements

Our work would not have been possible without the support of the many landowners who gave us permission to place cameras on their land, including K. Konrad, C. Konrad, W. Silva, M. Hinostroza, L. Miño H. Beckhert, K. Beckhert, V. Beckhert, and G. Weisser. We thank the Applied Wildlife Ecology Lab especially N.A. Arringdale for GIS support and K.L. Mills for feedback and edits on earlier drafts, and C. Badgley, A.J. Marshall, and J. Vandermeer for assistance with study design. We appreciate field support from M.A. Lyons in helping collect data and communicate in Spanish with landowners. In additional, this work would have been impossible without the support of M. Jiménez, who generously provided logistical support throughout the field season. Finally, we would like to thank the University of Michigan Latin American and Caribbean Association and the Rackham Graduate school for funds which made the data collection possible.

---

**Table 1.** Top occupancy models for every species (occupancy(Ψ), detection(p)). QAICc was used instead of AICc in model ranking for güiña occupancy to account for overdispersion of the global model.

| Species | Top models | AICc | ΔAICc* | $w_i$ |
| --- | --- | --- | --- | --- |
| Chilla | p(dogo, 10uds) $\Psi$(iso) | 429.809 | 0 | 0.175 |
| | p(dogo, 10uds, cam) $\Psi$(iso) | 430.289 | 0.480 | 0.138 |
| | p(dogo, 10uds) $\Psi$(dogo) | 430.613 | 0.804 | 0.117 |
| | p(dogo, 10uds) $\Psi$(dogo, 10uds) | 430.782 | 0.974 | 0.108 |
| | p(dogo, 10uds, cam) $\Psi$(dogo) | 430.948 | 1.139 | 0.099 |
| | p(dogo, 10uds) $\Psi$(iso, dogo) | 431.139 | 1.331 | 0.090 |
| | p(dogo, 10uds, cam) $\Psi$(dogo, 10uds) | 431.511 | 1.702 | 0.075 |
| | p(dogo, 10uds) $\Psi$(iso, 10uds) | 431.736 | 1.927 | 0.067 |
| | p(dogo, 10uds) $\Psi$(~1) | 431.738 | 1.929 | 0.067 |
| | p(dogo, 10uds) $\Psi$(iso, dogo, 10uds) | 431.792 | 1.982 | 0.065 |
| Culpeo | p(dogo, cam, 10uds) $\Psi$(dogo, sm) | 124.932 | 0 | 0.273 |
| | p(dogo, 10uds) $\Psi$(dogo, sm) | 126.238 | 1.306 | 0.142 |
| | p(dogo, cam, 10uds) $\Psi$( sm) | 126.280 | 1.348 | 0.139 |
| | p(dogo, cam) $\Psi$(dogo, sm) | 126.598 | 1.666 | 0.119 |
| | p(dogo, trap, 10uds) $\Psi$(sm) | 126.734 | 1.802 | 0.111 |
| | p(dogo, 10uds) $\Psi$(sm) | 126.753 | 1.821 | 0.110 |
| | p(dogo, cam, trap) $\Psi$(dogo, sm) | 126.802 | 1.870 | 0.107 |
| Dog | p(10uds, trap) $\Psi$(forest) | 272.000 | 0 | 0.370 |
| | p(10uds, trap, cam) $\Psi$(forest) | 272.306 | 0.306 | 0.318 |
| | p(10uds, trap) $\Psi$(forest, 10uds) | 273.704 | 1.704 | 0.158 |
| | p(10uds, trap, cam) $\Psi$(forest, 10uds) | 273.757 | 1.757 | 0.154 |
| Güiña | p(~1) $\Psi$(~1) | 6.830 | 0 | 0.101 |

## Supplementary Material

**Figure S1.**
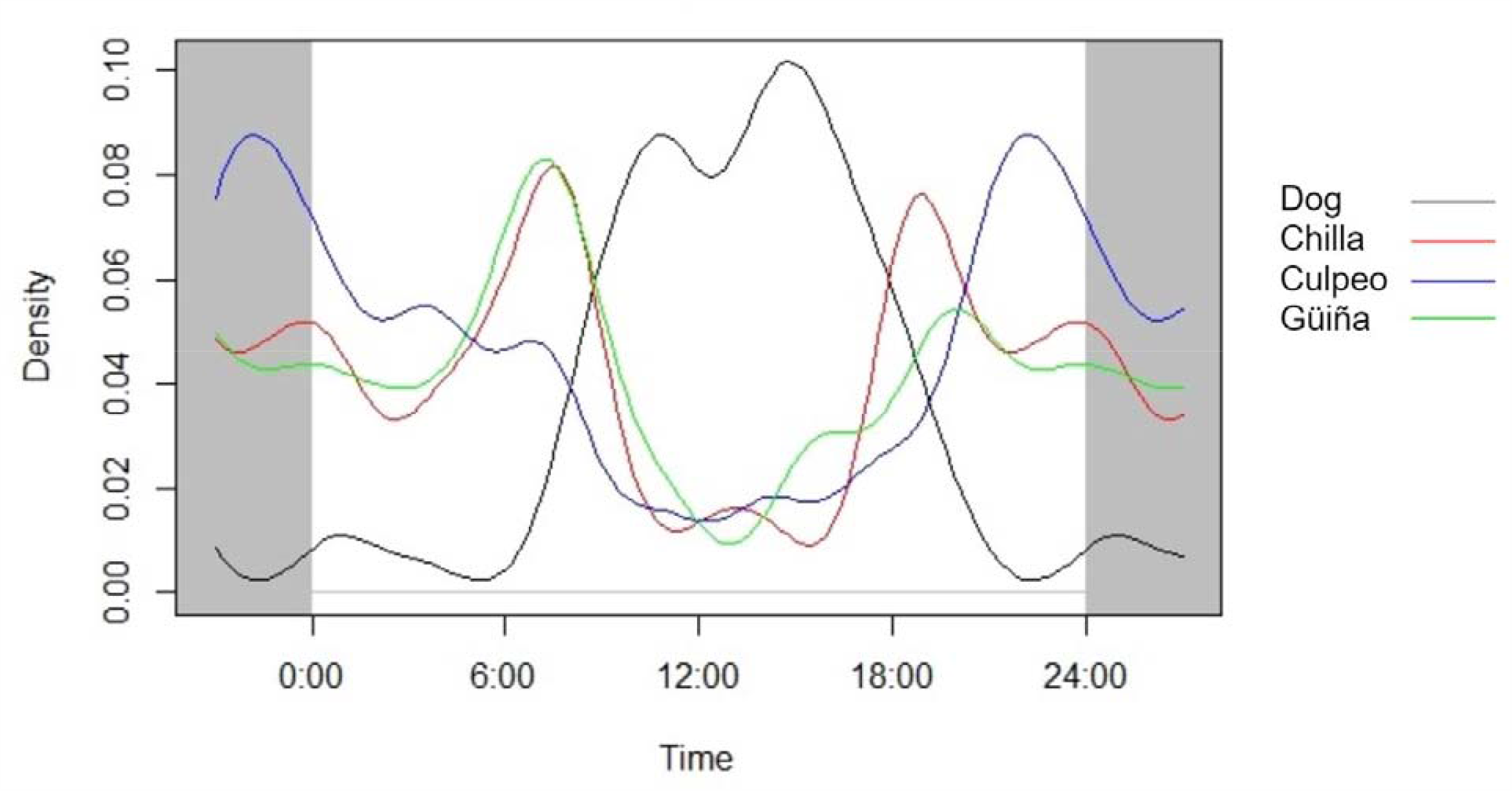
24-hour activity patterns of all four study species. Dogs are distinct from native species in having a clearly diurnal activity pattern.

**Figure S2.**
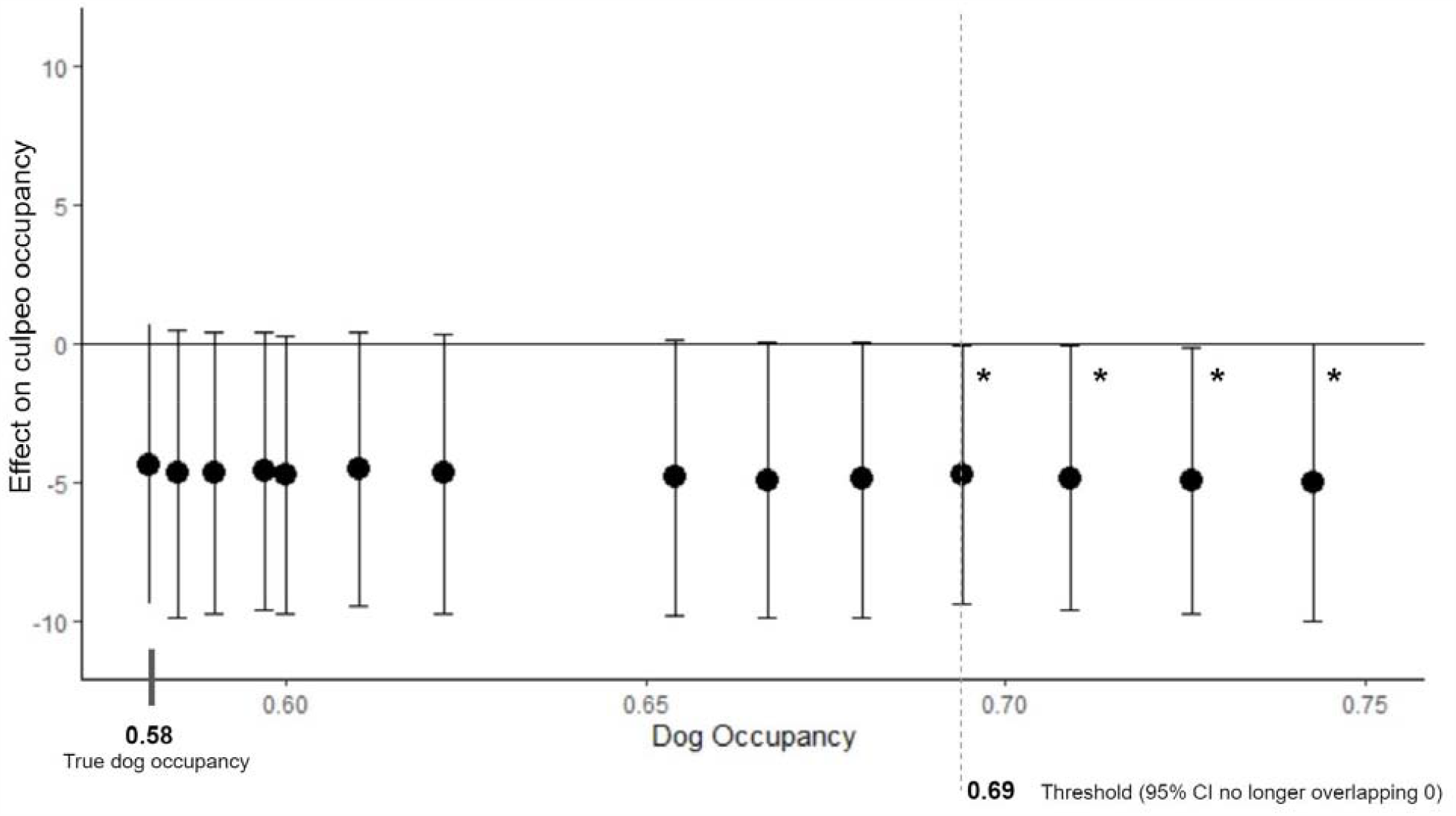
The effect of a simulated increase in dog occupancy across the landscape on the beta coefficient and 95% confidence interval for the *dogo* covariate in the culpeo occupancy model. *represents a significant beta coefficient.

**Table S1.** Overlap coefficients (Δ) and Mardia-Watson-Wheeler test for homogeneity of means for every pairwise combination of the study species. Δ4 was used for every comparison except for those pairs containing the culpeo, where Δ1 was used to account for lower number of triggers.

| Comparison | $\Delta$ Overlap (95% CI) | W statistic | *p*-value |
| --- | --- | --- | --- |
| Chilla-Dog | 0.40 (0.25-0.44) | 83.62 | <0.001 |
| Culpeo-Dog | 0.35 (0.22-0.48) | 42.96 | <0.001 |
| Guigna-Dog | 0.43 (0.30-0.56) | 47.84 | 0.001 |
| Chilla-Culpeo | 0.78 (0.66-0.89) | 5.02 | 0.08 |
| Chilla-Guina | 0.89 (0.80-0.97) | 0.47 | 0.79 |
| Guina-Culpeo | 0.78 (0.75-1.03) | 4.79 | 0.09 |

## Notes

### Competing Interest Statement

The authors have declared no competing interest.

